# Cellular and mitochondrial correlates of regional glucose metabolism in the human cortex

**DOI:** 10.64898/2026.10.22.760701

**Authors:** Matej Murgaš, Sebastian Habernig, Christian Milz, Samantha Graf, Murray B Reed, Benjamin Eggerstorfer, Pia Falb, Clemens Schmidt, Elisa Briem, Gabriel Schlosser, Alexandra Mayerweg, Kenan Al Barede, Ivan Pörnbacher, Lukas Nics, Sazan Rasul, Marcus Hacker, Dan Rujescu, Andreas Hahn, Rupert Lanzenberger

## Abstract

Regional differences in human brain glucose metabolism reflect local energy demands, but their cellular and mitochondrial determinants remain unclear. We combined [^18^F]FDG positron emission tomography in healthy human participants with estimates of cellular composition and mitochondrial density to unravel (sub)cellular correlates of heterogeneity in brain glucose metabolism. By correcting for RNA content and differential gene expression due to postmortem sampling, we addressed methodological shortcomings in multimodal neuroimaging integration. We introduce transcriptionally derived cytoarchitectonic maps of the human cortex that are consistent with histological findings, providing a biologically grounded framework for relating *in vivo* neuroimaging signals. Accordingly, mitochondrial distribution showed strong association with the intrinsic energy requirements at resting state. In contrast, relations between metabolism and neuronal composition were driven by sensory input and mitochondria-sparse thin-axon neurons, suggesting that stimulation-induced energy demand is met rather by neuronal processes that are independent of mitochondrial abundance to maintain efficient information processing.

## Introduction

The human brain has a high energy demand relative to its size ^1–4^, driven by the metabolic costs of neuronal network activity, synaptic transmission, and maintenance of ionic gradients ^5, 6^. Although multiple interacting metabolic pathways contribute to the human brain’s energy landscape ^7^, glucose remains its primary energy substrate ^1, 8^. A standard approach to measuring regional cerebral glucose metabolism (CMRGlu) *in vivo* is [^18^F]fluorodeoxyglucose ([^18^F]FDG) PET ^9, 10^, which indirectly reflects neuronal and synaptic activity ^5, 6, 11^. However, despite ongoing methodological advancements in [^18^F]FDG PET ^12–14^, the signal remains a macroscopic measure of multiple cellular and metabolic processes within the brain. For example, glial cells such as microglia and astrocytes affect the [^18^F]FDG PET signal under certain functional and metabolic conditions ^15–17^. Acknowledging these complexities, the integration of diverse datasets enables multiscale approaches that provide new opportunities to unravel composite signals, such as those derived from neuroimaging ^18, 19^.

Mitochondria contribute an additional functional layer by engaging in complex biological processes, thereby playing a crucial role in energy supply ^20^. These processes include cognitive function ^21, 22^, modulation of certain psychiatric phenotypes, signal transduction ^23^, regulation of neuronal activity dynamics ^24^ and neurotransmitter release ^25^. These are facilitated through its central function of generating adenosine triphosphate (ATP) through oxidative phosphorylation, a core element of the brain’s molecular energetic landscape ^26, 27^. Spatial and morphological characteristics of these organelles likely shape regional metabolic differences and the overall energy-transformation capacity of certain brain regions ^27, 28^, potentially enabling differential interpretation of the [^18^F]FDG signal. Moreover, alterations in mitochondrial function have been associated with psychiatric conditions ^29, 30^ and neuroinflammation in neurodegenerative disorders_31, 32._

Resources, such as the Allan Human Brain Atlas, provide spatially anchored genome-wide expression profiles and enable the interpretation of other brain phenotypes through the lens of single-gene expression data, cell-type abundances and biological pathways ^19, 33, 34^. Thus, an additional layer worth investigating involves the use of marker genes to derive transcriptomic “fingerprints” of specific cell types, which might help to uncover the underlying cellular architecture of the human brain ^35, 36^. Regional heterogeneity of the human brain’s cellular architecture hints at varying metabolic requirements to reflect the cellular mosaic that constitutes each region ^35, 37^. Integrating cell-type architecture might reveal new insights into regional mitochondrial and [^18^F]FDG signals, further illuminating the multiscale complexity of the human brain ^19^.

Neuroscience research progressively employs multimodal and multiscale approaches that leverage multiomic data to investigate the human brain ^18, 19, 38, 39^. [^18^F]FDG PET examines glucose metabolism *in vivo* ^40^, enzymatic abundances of mitochondrial content reveal the molecular energetic landscape ^27^, and transcriptomic atlases illuminate the cell-type composition of the human brain ^35, 41–43^. However, these multimodal brain phenotypes have largely been addressed separately, which limits inferences about how metabolic, molecular, and cellular features relate to one another. In the present study, we aimed to create comprehensive voxel-wise representations of relative cellular proportions in the human cerebral cortex to facilitate the integration of cytoarchitecture into *in vivo* neuroimaging studies. To this end, we adopted an integrative approach that aims to complement the shortcomings of each modality.

## Results

### Voxel-wise relative cell type proportions match histological data

Cell type distributions cannot be measured directly in the living human brain, so we estimate them from the expression of cell type-specific marker genes. The Allen Human Brain Atlas ^34^, however, provides microarray expression only at a limited number of discrete tissue samples, whereas our typical neuroimaging data are available at much higher spatial resolution. To integrate the cell type distributions with diverse neuroimaging phenotypes, we first generated voxelwise maps of marker gene expression by processing the AHBA microarray data in multiple steps. (Fig. 1a).

**Figure 1.**
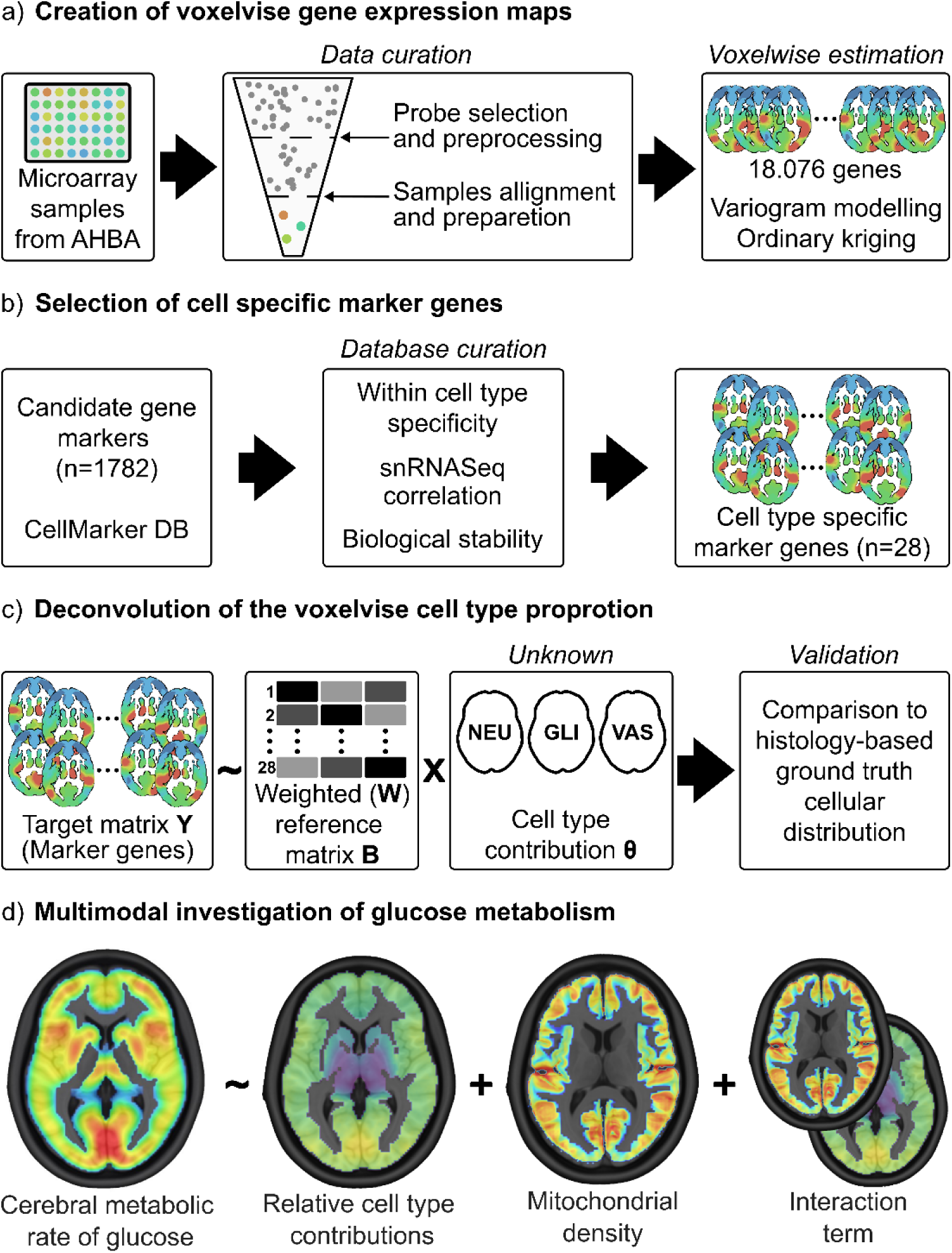
Overview of the relative cell-type distribution estimation and the analysis. a) Voxel-wise estimation of whole-brain gene expression patterns yields 18076 brain maps. b) 28 cell-type-specific marker genes were selected from the curated marker database, CellMarker ^47^. c) Using ridge regression, neuronal, glial, and vascular relative cell-type proportions were estimated. d) Integrating cortical maps of mitochondrial density and relative cell-type proportions into one model, we investigate their impact on the cerebral glucose metabolism rate.

First, probes were filtered based on interprobe correlation, spatial variability, and proximity to snRNASeq signal. After probe filtering, we retained 38,595 probes across 18,076 genes for further analysis. After quality control and anatomical assignment, samples were mirrored along the X axis, yielding 7,248 mirrored samples covering the cortex, subcortex, cerebellum, and brainstem. Spatial interpolation using variogram modelling and ordinary kriging yielded voxel-wise gene expression maps for all selected markers. Here we have refined an established approach, previously published by our group ^44^. Specifically, we used re-evaluated MNI coordinates ^45^, incorporated concordance between microarray data and snRNASeq signals ^42^ into probe selection, and extended the original kriging space to encompass a 10% thresholded grey matter map from Harvard-Oxford as implemented in FSL ^46^. The resulting mRNA expression maps demonstrate distinct spatial patterns across major brain regions. The database of all 18,076 spatial gene expression patterns is available online. The presented spatial gene expression patterns served as input for deconvolution of cellular contributions.

Cell-specific candidate marker genes of neuronal, glial and vascular cells were compiled from a curated database ^47^ and refined using cell-type specificity derived from single-nucleus RNA sequencing (snRNASeq) ^42^ (Fig. 1b). From an initial set of 1,173 candidate markers, 28 genes were retained following specificity filtering and redundancy reduction (see Methods). Selected markers showed high specificity (specificity score > 2.0) within their target cell types relative to competing cell classes. The final marker set comprised ten marker genes for neurons, nine for glial cells and nine for vascular cells. To facilitate interpretation, marker genes for each cell type are shown in Fig. 2a-c (Suppl. Table 1), illustrating canonical expression patterns.

**Figure 2.**
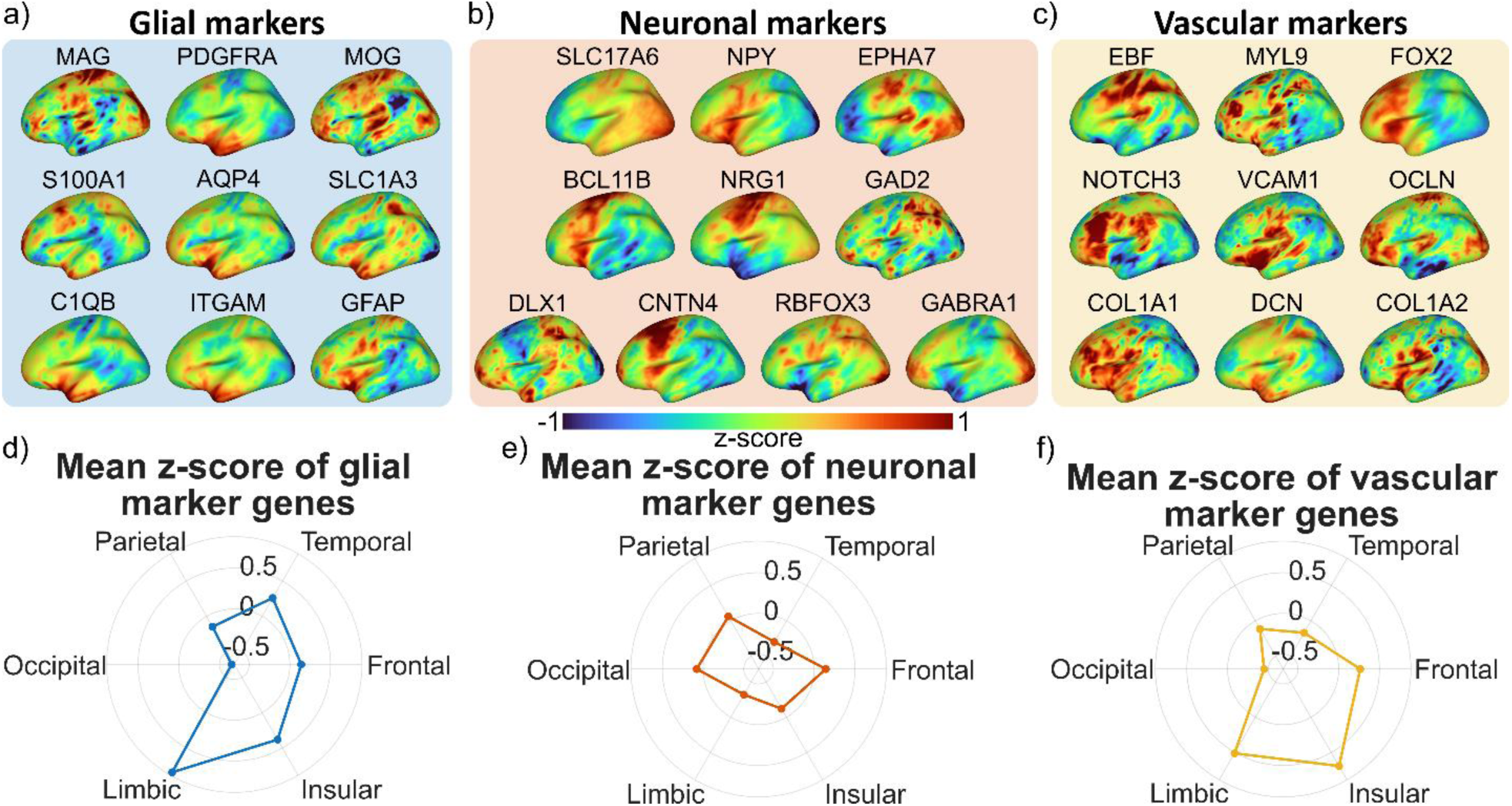
Definition of the set of marker genes. Surface representation of marker genes used to estimate relative a) glial, b) neuronal and c) vascular cell proportions. Spider plots showing distinct mean regional z-scored values of d) glial cell marker genes, e) neuronal cell marker genes and f) vascular cell marker genes. Glial cell marker genes exhibited the highest average z-scored values in frontal, temporal, insular and limbic regions. Parietal, occipital and frontal regions exhibited the highest mean z-scored values of the neuronal gene set of markers, while the vascular gene set displayed the highest expression patterns in insular and limbic regions.

To summarise the spatial heterogeneity of selected cell-specific marker genes, mean z-scored expression values were calculated across major cortical regions (frontal, temporal, parietal, occipital, limbic, and insular cortex). The resulting marker-derived expression patterns revealed distinct regional variation across cellular classes. Within glial marker sets (Fig. 2a), the frontal, temporal, limbic and insular cortices showed the highest expression (Fig. 2d), whereas neuronal marker sets (Fig. 2b) demonstrated increased expression in the frontal, parietal and occipital cortices (Fig. 2e). The spatial transcriptomic patterns of these two cell classes (Fig. 2d-e) are in line with the previously reported glia-to-neuron ratios showing a decrease in the anterior-to-posterior direction ^48^. Vascular marker sets showed higher expression in insular and limbic regions than in other cortical areas (Fig. 2f). Utilised mRNA expression patterns provide a heterogeneous spatial characterisation of the cell-type-associated marker sets, consistent with previous literature ^48–50^, and are used for subsequent deconvolution.

Thereafter, the cell type-specific voxelwise marker gene expression maps were entered into a non-negative ridge regression ^51^ to estimate the relative proportions of three complementary cellular classes (Fig. 1c): relative glial proportion (GLI_R_) (Fig. 3a), relative neuronal proportion (NEU_R_) (Fig. 3b), and relative vascular proportion (VAS_R_) (Fig. 3c) across the cortex. The model accounts for the proximity between snRNASeq ^42^ and microarray signal ^34^, as well as a biological stability metric ^52^, that represents expression invariance between the *in vivo* and *post-mortem* states. Additionally, RNA-content correction accounted for differences in total transcript abundance among cell classes, yielding estimates of relative cellular contributions. Consistent with previous transcriptomic studies ^42, 53^, our continuous spatial maps demonstrated distinct patterns of cellular organisation across the cortex. Glial cells showed the highest GLI_R_ in the prefrontal, insular, parietal and temporal cortices (Fig. 3a, d) and the lowest in the occipital and primary visual cortices (V1). In contrast, NEU_R_ was highest in the primary visual cortex (Fig. 3b, e). Vascular cells showed the lowest VASR across the cortex but contributed most in the insular and cingulate cortices (Fig. 3c, f). The ratio between estimated GLI_R_ and VAS_R_ (23% - 25%) aligns with previously reported results of 30% non-neuronal cells being non-glial cells ^50^. The voxel-wise NEU_R_, GLI_R_ and VAS_R_ maps are publicly available.

**Figure 3.**
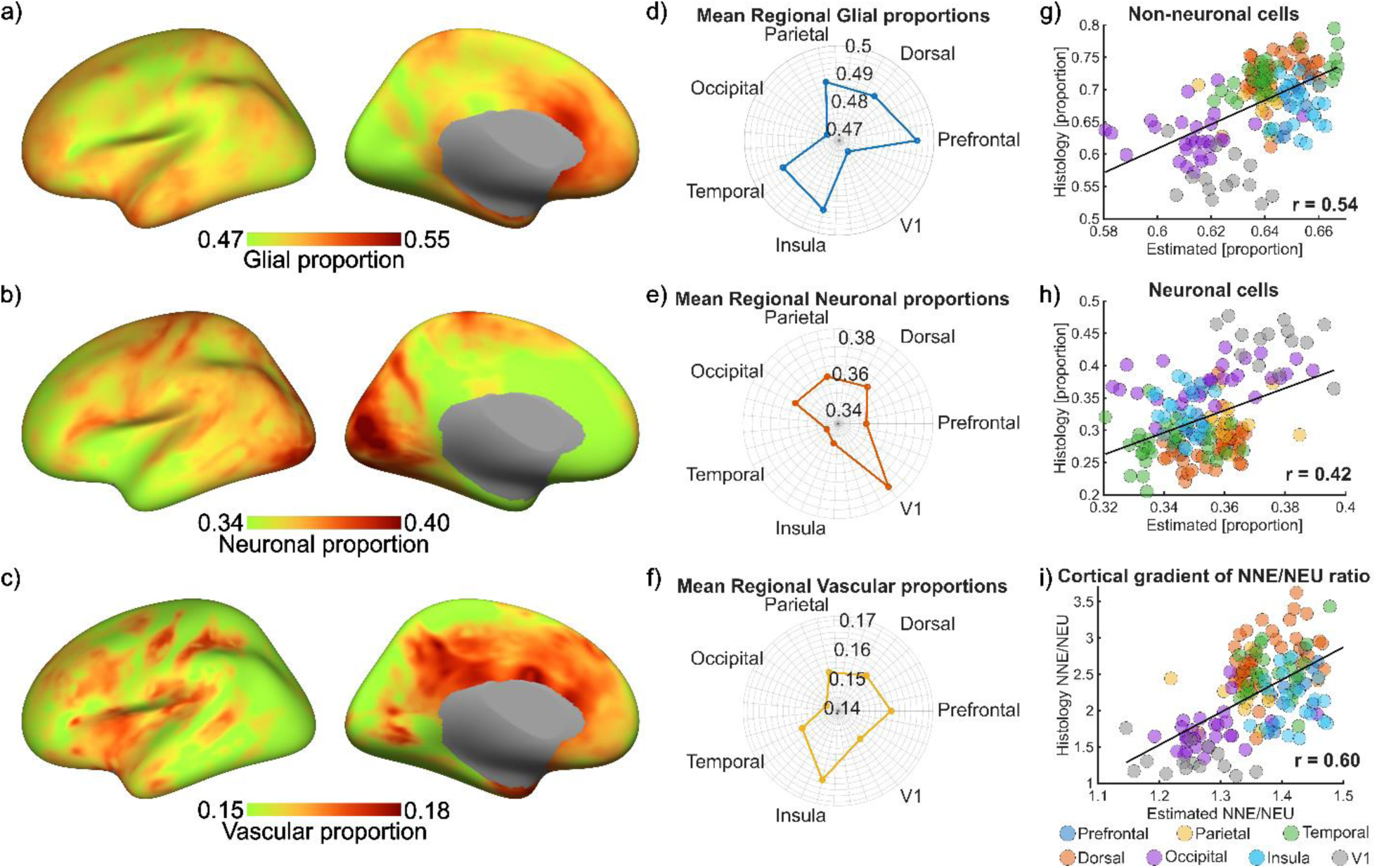
Spatially continuous representation of cellular proportions. Cortical projection of a) relative glial proportion, b) relative neuronal proportion and c) relative vascular proportion estimated using non-negative ridge regression. Spider plots showing the mean regional relative proportions of d) glial cells, e) neuronal cells, and f) vascular cells. Comparison of estimated and histology-based regional proportions of g) non-neuronal and h) neuronal cell types, including the i) non-neuronal-to-neuronal (NNE/NEU) ratio. The regions are based on the original histological data published by Ribeiro et al.^48^.

The biological validity of the cell-type contribution maps was further assessed by comparing them with previously reported ground-truth cellular proportions ^48, 50, 54^. Specifically, estimated neuronal and non-neuronal contributions (NNE), representing the combined relative proportions of glial and vascular cells, were compared with the histology-based proportions. A good match was observed for the relative contribution of non-neuronal cells (Fig. 3g, r=0.54) and for relative neuronal proportions (Fig. 3h, r=0.42). Comparison of the cortical gradient regarding the non-neuronal-to-neuronal cell ratio (NNE/NEU) showed even stronger agreement between estimated and histology-based data (Fig. 3i, r=0.60). Together, these results suggest that the estimated maps may provide a reliable spatial proxy for regional variation in cellular composition and, hence, foster the investigation of cell-type-associated spatial patterns in neuroimaging studies.

### Baseline glucose metabolism is modulated by mitochondrial density and glia-to-neuron ratio

To investigate the involvement of the obtained cellular composition and mitochondrial density (MitoD) in baseline glucose metabolism, we have integrated our estimated relative cellular proportions into a weighted linear regression model (Fig. 1d). We found that regional CMRGlu measured with eyes closed (CMRGlu_EC_) was positively associated with mitochondrial density (Fig. 4b; t(101)=4.06, p_spin_<0.05, r=0.39). The relationship between glia-to-neuron (G2N) ratio and CMRGlu_EC_ was also positive but not significant (Fig. 4a; t(101)=4.86, p_spin_=0.11, r=0.32). The interaction effect between MitoD and G2N showed a significant association with CMRGlu (t(101)=3.13, p_spin_<0.05). The Johnson-Neyman plot (Fig. 4c) indicates that the positive association between MitoD and CMRGlu_EC_ was significant specifically in regions with a higher glia-to-neuron ratio. In contrast, the association between the glia-to-neuron ratio and CMRGlu was significant specifically in regions with higher mitochondrial distribution.

**Figure 4.**
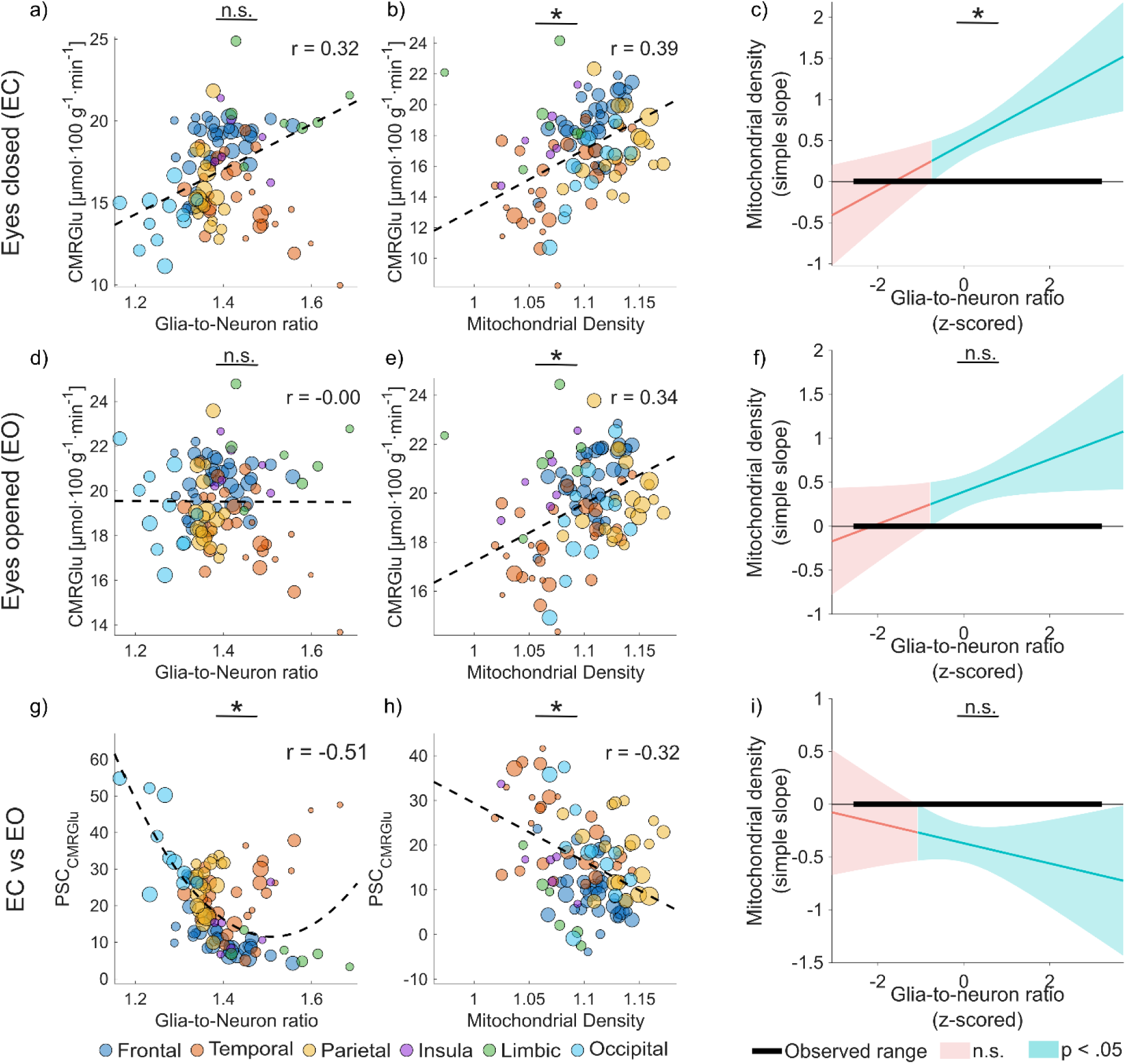
Biological model integrating baseline glucose metabolism, mitochondrial density and glia-to-neuron ratio. The size of the marker represents the size of the regions used in weighted linear regression as a scaling factor, and the colour represents the marker association with gross cortical regions - frontal (dark blue), temporal (red), parietal (yellow), insular (violet), limbic (green), and occipital (cyan) lobes. Positive associations were observed between resting-state CMRGlu during eyes-closed conditions (CMRGlu_EC_) and a) the G2N ratio and b) mitochondrial density (MitoD). d) No association of the G2N ratio with CMRGlu during the eyes-open (CMRGlu_EO_) state was shown, while f) mitochondrial density exhibited a negative effect on CMRGlu_EO_. Association of the MitoD and G2N ratio with the per cent signal change (PSC_CMRGlu_ = [CMRGlu_EO_/CMRGlu_EC_ - 1] × 100) is shown in panels g) and h), respectively. The interaction effects between the G2N ratio and MitoD during c) eyes closed, f) eyes opened and i) PSC_CMRGlu_ are represented by Johnson-Neyman plots. These plots show the conditional slopes of MitoD as a function of the G2N ratio. The model, including a quadratic term, shows the total slope of the G2N ratio as a function of MitoD, accounting for non-linearity. Solid lines represent the model-estimated simple slope at each value of the moderator; shaded bands represent the corresponding 95% confidence interval. Teal shading indicates values of the moderator at which the simple slope is statistically significant (p<0.05); pink shading indicates the slope is not significantly different from zero. The thick black horizontal bar denotes the observed range of the moderator variable in the sample. Statistically significant post-hoc tests are marked with * and non-significant as n.s. The dependent variable is always corrected for the respective covariates that are not included in the post hoc comparisons and scatter plots.

To further emphasise these associations and elucidate the involvement of individual cell types, we replace the glia-to-neuron ratio with separate relative neuronal and glial proportions. NEU_R_ was negatively associated with CMRGlu_EC_ (Suppl. Fig. 1a, t(101)=-5.34, p_spin_ <0.05, r=-0.41), whereas MitoD remained positively associated with CMRGlu_EC_ (Suppl. Fig. 1b, t(101)=5.52, p_spin_ <0.05, r=0.43). Moreover, the significant interaction effect between NEU_R_ and MitoD (t(100)=-3.70, p_spin_<0.05), together with the Johnson-Neyman plot (Suppl. Fig. 1c), indicates that the association between mitochondrial distribution and glucose metabolism depends on the relative neuronal composition of cortical regions. In contrast, GLI_R_ was not significantly associated with CMRGlu_EC_ after accounting for MitoD (Suppl. Fig. 2a-c). Overall, these results indicate that the G2N ratio likely reflects variation in neuronal proportions rather than an isolated independent link between glial proportion and CMRGlu_EC_. This aligns with the distinct metabolic characteristics of neurons and glia, i.e. their individual reliance on glycolytic and mitochondrial oxidative pathways ^1, 55^.

### Sensory input affects glucose metabolism independent of mitochondrial density

Next, we examined if the observed associations between glucose metabolism, MitoD and cellular composition depend on the sensory input, using a model in which CMRGlu was measured with eyes open (CMRGlu_EO_). The positive relationship with MitoD was preserved (Fig. 4e, t(101)=4.17, p_spin_<0.05, r=0.34); however, the effect of the G2N on CMRGlu_EO_ was strikingly absent (Fig. 4d, t(101)=-0.04, p_spin_=0.99, r=-0.00). In addition, the Johnson-Neyman decomposition shows that the conditional association between MitoD and CMRGlu is highly sensitive to the G2N and lacks consistent statistical support across the observed data range, consistent with the interaction term not being significant (Fig. 4f, t(101)=2.05, p_spin_=0.10). Splitting the analysis by cell type showed a significant interaction effect between NEU_R_ and MitoD with CMRGlu_EO_ (t(101)=-2.56, p_spin_<0.05). No other significant effect of singular cellular proportions (Suppl. Fig. 1d-f, Suppl. Fig. 2d-f) was observed. These results suggest that mitochondrial abundance is a key determinant shaping regional glucose metabolism in general, whereas the relationship between neuronal composition and glucose metabolism depends on the functional condition ^56^.

When examining the relative change in glucose metabolism between eyes-open and eyes-closed conditions, the percent signal change of CMRGlu (PSC_CMRGlu_) was significantly associated with the glia-to-neuron ratio (Fig. 4g, t(100)=-6.70, p_spin_<0.05, r=-0.51) and with mitochondrial density (Fig. 4h, t(100)=-4.10, p_spin_<0.05, r=-0.32). Exploratory analyses replacing G2N with NEU_R_ and GLI_R_ showed a strongly significant association between relative neuronal proportion and its quadratic term with PSC_CMRGlu_ (Suppl. Fig. 1g, Suppl. Table 2). Conversely, GLI_R_ was negatively associated with PSC_CMRGlu_ (Suppl. Fig. 2g, Suppl. Table 3). The model including NEU_R_ showed a significant negative association between MitoD and PSC_CMRGlu_ (Suppl. Fig. 1h, Suppl. Table 2), whereas the model including GLI_R_ showed a non-significant negative association (Suppl. Fig. 2h, Suppl. Table 3).

Disentangling the analysis to individual cortical regions unveiled diverse associations between PSC_CMRGlu_, MitoD and relative cellular proportions (Suppl. Fig. 3). Focusing on the occipital lobe as the primary cortical region for visual sensory input, we observed a similar association between relative cellular proportions, mitochondrial density, and PSC_CMRGlu_ (Suppl. Fig. 3). Interestingly, the same associations were preserved in the frontal lobe, which contributes to visual attention and top-down modulation of sensory processing ^57^. In contrast, the parietal, temporal, insular and limbic lobes showed association with PSC_CMRGlu_ in the opposite direction. These results may reflect a heterogeneous relationship between molecular organisation of cortical regions and metabolic adaptation during sensory input-induced transition.

### Transcriptional signatures of axonal diameter reflect sensory input-dependent glucose metabolism

Previous work has proposed that during periods of peak energy demand, the human brain may prioritise information efficiency over energy efficiency ^56^. It was suggested that thin axons (< 0.5 μm), exhibiting limited numbers of mitochondria due to their biophysics ^58^, meet their additional energetic demands through aerobic glycolysis, shifting their ATP production away from the more energy-efficient oxidative phosphorylation ^56^. To further elucidate the role of cellular composition and mitochondrial density in sensory input-induced alterations in CMRGlu (PSC_CMRGlu_ defined above), we included expression patterns of marker genes for thin-(CUX1 and CUX2) and thick-axon (FEZF2 and BCL11B) neurons in the weighted linear regression. This allowed us to evaluate potential associations of transcriptional signatures related to neuronal populations of axonal calibre on PSC_CMRGlu_.

Our findings suggest that marker genes of thin- and thick-axon neurons exhibit opposing spatial signatures in their relationships with mitochondrial density (MitoD) and stimulation-related glucose utilisation. Across cortical regions, CUX1 (Fig. 5a) and CUX2 (Fig. 5b) expression levels showed a positive association with the PSC_CMRGlu_ (t(100)=4.78, p_spin_<0.05, r=0.71) and t(100)=7.24, p_spin_<0.05, r=0.64), respectively. In contrast, FEZF2 (Fig. 5c) and BCL11B (Fig. 5d) expression levels showed a negative, however non-significant association (t(100)=-4.13, p_spin_=0.11, r=-0.32) and t(100)=-4.92, p_spin_=0.10, r=-0.32). In all cases, mitochondrial density showed a negative association with PSC_CMRGlu_, which was however only significant for CUX1 (Fig. 5a right column, t(100)=-3.86, p_spin_<0.05, r=-0.33) and CUX2 (Fig. 5b right column, t(100)=-4.72, p_spin_<0.05, r=- 0.39) and no significant interaction effects.

**Figure 5:**
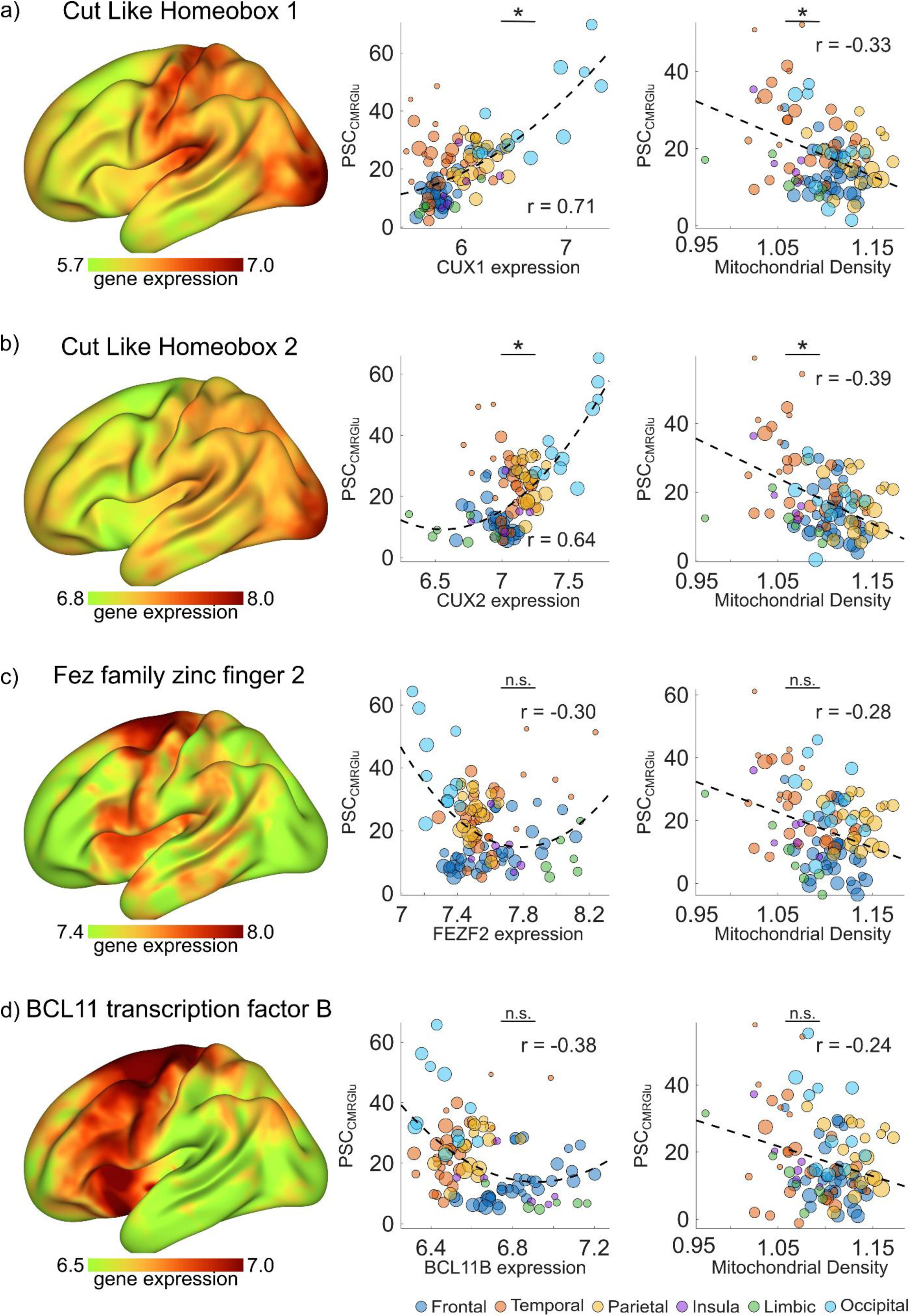
Integration of marker genes for neurons with thin and thick axonal diameter. a) Cortical representation of the Cut-like Homeobox 1 (CUX1), the marker gene for thin axonal neurons (left). Model investigating CUX1 and mitochondrial density (MitoD) on per cent signal change in cerebral metabolic rate of glucose (PSC_CMRGlu_). b) Cortical representation of the Cut-like Homeobox 2 (CUX2), the marker gene for thin axonal neurons (left). Model investigating CUX2 and MitoD on PSC_CMRGlu_. c) Cortical representation of the Fez family zinc finger 2 (FEZF2) the marker gene for thick axonal neurons (left). Model investigating FEZF2 and MitoD on PSC_CMRGlu_. d) Cortical representation of the BCL11 transcription factor B (BCL11B), the marker gene for thick axonal neurons (left). Model investigating BCL11Band MitoD on PSC_CMRGlu_. Statistically significant post-hoc test are marked with <u>*</u> and non-significant as <u>n.s..</u>

Whole-cortex associations might partly reflect a regional correspondence between gene expression signatures and sensory input-specific recruitment of the individual lobes. However, the same directional pattern was observed when the analyses were limited to the visual cortex as the primary target region of the visual input. Within the visual cortex, CUX1 and CUX2 expression levels showed a negative association with MitoD (CUX1: Suppl. Fig. 4a, r=-0.30; CUX2: Suppl. Fig. 4b, r=-0.47), and remained positively associated with PSC_CMRGlu_ (CUX1: Suppl. Fig. 4a, r=0.62; CUX2: Suppl. Fig. 4b, r=0.67). In contrast, FEZF2 and BCL11B expression levels showed a positive or no correlation with MitoD (FEZF2: Suppl. Fig. 4c, r=0.36; BCL11B: Suppl. Fig. 4d, r=0.01) and a negative association with PSC_CRMGlu_ (FEZF2: Suppl. Fig. 4c, r=-0.36; BCL11B: Suppl. Fig. 4d, r=-0.44). The bidirectional relationships of these transcriptional profiles align with the efficiency-tradeoff hypothesis ^56^, supporting the notion that aerobic glycolysis contributes more upon stimulation in mitochondria-sparse thin-axon neurons during periods of rapidly shifting energy demands.

## Discussion

The metabolic demands of the human brain arise from the coordinated activity of diverse cell populations and their underlying molecular mechanisms. Integrating regional glucose metabolism with cellular composition and mitochondrial abundance may therefore help to characterise the cellular and molecular features underlying differences in cerebral energy requirements. Using a new version of the approach established by our group ^44^, we generated enhanced whole-brain voxel-level gene expression patterns. Subsequently, we deconvolved these transcriptional signatures to estimate the spatially resolved relative proportions of major cell types across the cortex. Previous works estimated cell-type distribution at the region-^43^ and voxel-wise levels ^59^ from transcriptomic signatures. However, these estimates reflect RNA contributions rather than cellular abundance, as cell types differ in RNA content ^60^. Here, we used snRNAseq-derived cell-type profiles from Siletti et al. ^42^ and incorporated cell-type-specific RNA content into the deconvolution to adjust the RNA-based estimates to reflect relative cellular abundance. In addition, we used estimates of postmortem-related transcriptional changes reported by Liharska et al.^52^ to reduce the influence of postmortem-related variation in gene expression.

The resulting relative cellular contribution maps showed spatial patterns consistent with previously reported histological distributions of neuronal and non-neuronal cells. Within the cortex, we observed a pronounced anterior-posterior gradient, with neuronal proportions increasing toward posterior regions ^48^. Relative to frontal regions, visual cortical areas exhibited particularly high neuronal density, consistent with previous histological observations in the human visual cortex ^48, 61^. This pattern is consistent with established regional variation in cortical cytoarchitecture, particularly the notion that neuronal density varies systematically across cortical areas rather than scaling linearly with grey-matter volume ^48, 54, 62^. Correspondingly, the glia-to-neuron ratio decreased along the rostral-caudal axis, reflecting regionally varying cellular composition across the cortex ^48, 50^. In line with these observations, higher glia-to-neuron ratios have been reported in frontal cortical regions, including the prefrontal cortex, compared to other cortical areas ^49, 50^, further highlighting the progressive shift toward greater neuronal contribution in posterior cortical regions. These histological comparisons support the biological validity of our deconvolution-derived estimates by showing that they capture meaningful spatial variation in cortical cellular composition. Consequently, these spatially continuous estimates of relative neuronal, glial, and vascular proportions may facilitate the direct investigation of how regional cellular composition relates to *in vivo* neuroimaging and imaging transcriptomic studies.

The human brain is an energetically demanding organ that relies primarily on glucose to sustain neuronal signalling ^63^, with approximately 25-40% of the brain’s glucose consumption attributed to housekeeping and maintenance of the resting potential ^5, 64^. Using CMRGlu during eyes-closed as a proxy for non-stimulation-related energy demand, we showed a positive association between MitoD and CMRGlu_EC_, whereas relative neuronal distribution was negatively associated with CMRGlu_EC_. This relationship was further supported by a significant interaction between mitochondrial density and relative neuronal distribution, showing that the association between CMRGlu_EC_ and MitoD was stronger in neuronal-low regions. This result is consistent with mitochondrial housekeeping demands contributing more to baseline metabolism in regions with a lower relative neuronal proportion. These regions are typically characterised by fewer neuronal cell bodies and a greater abundance of neuropil and supporting glial elements ^65^. Consequently, stimulation-related neuronal metabolism may contribute less in relation to total glucose metabolism in these regions. However, this interpretation is limited by the lack of neuronal and glial mitochondrial markers.

Eye opening shifts the visual cortex from an intrinsic state to sensory input processing ^66^, which is increased neuronal engagement as reflected in a proportionally higher glucose utilisation relative to the eyes-closed condition. Indeed, the relative neuronal proportion showed a positive nonlinear association with PSC_CRMGlu_. Higher neuronal density likely reflects increased synaptic density and, thus, elevated glucose demand upon stimulation, consistent with prior evidence linking synaptic density to the metabolic rate of glucose ^37^. These observations are consistent with rodent data showing non-linear scaling of synaptic complexity in neuron-rich regions ^67^. On the other hand, upon stimulation, MitoD showed a negative association with PSC_CMRGlu_. Given the consistent negative association between PSC_CMRGlu_ and mitochondrial density in both eyes-closed and eyes-open conditions, it likely does not reflect reduced mitochondrial involvement during sensory-driven activation. It may instead indicate that mitochondria-dense regions already maintain higher baseline metabolic demand, resulting in a smaller relative increase in CMRGlu during activation ^1,68^.

The “efficiency tradeoff hypothesis” ^56^ proposes that during periods of peak energy demand, the human brain may prioritise information efficiency over energy efficiency. Therefore, thin axons (< 0.5 μm), which contain limited mitochondria due to their biophysics ^58^, may meet their additional energy demands through aerobic glycolysis, shifting their ATP production away from the more energy-efficient oxidative phosphorylation but toward rapid information processing at the cost of lower ATP yield per glucose molecule ^56^. Likewise, evidence in rodents and humans points towards recruitment of aerobic glycolysis during neuronal stimulation under varying conditions ^69–72^. Building on that hypothesis, we selected marker genes to spatially isolate putative thin- and thick-axon neuronal clusters within the human cortex. In rodents, CUX1 and CUX2 are established transcriptional markers for upper-layer II–IV pyramidal-neuron populations ^73, 74^, which include layer II/III intratelencephalic corticocortical and callosal projection neurons. Classical developmental studies provided the basis for identifying BCL11B ^75^ and FEZF2 ^76^ as potential candidate marker genes for thick-axon neuronal compartments. BCL11B plays a crucial role in axonal growth and connectivity ^75^, while FEZF2 represents an essential part during specification ^76^ of layer-5 extratelencephalic and other subcerebral projection neurons. In line with the efficiency tradeoff hypothesis, we found that thin-axonal neuronal marker genes were positively associated with sensory input-related changes in the cerebral metabolic rate of glucose, consistent with higher energetic demand, induced by the transition between functional conditions. In contrast, markers of thick-axonal neurons were negatively associated with PSC_CMRGlu_. These associations remained consistent in the occipital cortex, the target region of the sensory input-induced change. Notably, mitochondrial density was negatively associated with PSC_CMRGlu_ regardless of neuronal axonal diameter. Given mitochondria’s role in oxidative metabolism, the negative association may reflect differences in how regional mitochondrial abundance relates to condition-related changes in glucose metabolism. However, mitochondrial density alone does not determine oxidative capacity^1^.

While marker genes provide an excellent proxy for molecular distributions and properties, they should be interpreted cautiously. In particular, none of the markers discussed above directly measure axonal calibre, and although marker genes may be enriched in specific thin- or thick-axon architectures, considerable heterogeneity remains among individual neurons ^77–79^. Moreover, while microarray samples from the Allen Human Brain Atlas represent a valuable resource for gene expression data, they have several known limitations, such as imbalanced sampling sites, sex distribution, and longer post-mortem interval ^80^. Lastly, while the Human Brain Cell Atlas ^42^ provides RNA-seq data for approximately 3,000,000 cells, it does not currently include the frontal lobe.

This study revealed the involvement of cellular subtypes and mitochondria in glucose metabolism, demonstrating substantial mitochondrial involvement in housekeeping metabolism. Consistent with the efficiency tradeoff hypothesis, the association of thin-axonal neuronal markers with functional condition-related changes in CMRGlu suggests greater dependence on aerobic glycolysis under higher energetic demand. Overall, this study provides a framework for integrating spatially continuous cell-type estimates into neuroimaging, offering an additional perspective on the cellular basis of spatial variation in neuroimaging signals.

## Acknowledgments

We thank the graduated team members and the diploma students of the Neuroimaging Lab (NIL, head R. Lanzenberger) as well as the clinical colleagues from the Department of Psychiatry and Psychotherapy for clinical and/or administrative support, especially Gregor Dörl, Peter Stöhrmann, Lukas Artmeier, Vivienne Matev, Aurelia Sahl, Gregor Gryglewski, and Lisa Seilinger. The scientific project was performed with the support of the Research Platform Medical Imaging of the Medical University of Vienna.

## Author contribution statement (CRediT)

**Matej Murgaš**: Investigation, Conceptualisation, Methodology, Software, Validation, Formal analysis, Data Curation, Writing - Original Draft, Visualisation. **Sebastian Habernig**: Investigation, Writing - Original Draft, Visualisation. **Christian Milz**: Investigation, Data Curation. **Samantha Graf**: Investigation. **Murray B Reed**: Formal Analysis, Visualisation. **Benjamin Eggerstorfer**: Investigation. **Pia Falb**: Investigation. **Clemens Schmidt**: Investigation. **Elisa Briem**: Investigation. **Gabriel Schlosser**: Investigation. **Alexandra Mayerweg**: Investigation **Kenan Al Barede**: Investigation. **Ivan Pörnbacher**: Investigation. **Lukas Nics**: Supervision, Project administration. **Sazan Rasul**: Supervision. **Marcus Hacker**: Supervision, Resources. **Dan Rujescu**: Supervision, Resources. **Andreas Hahn**: Conceptualisation, Supervision, Resources, Funding acquisition. **Rupert Lanzenberger**: Conceptualisation, Visualisation, Supervision, Resources, Project administration, Funding acquisition. **All authors**: Writing - Review & Editing.

## Ethical considerations

The study was approved by the Ethics Committee of the Medical University of Vienna (ethics number: 1642/2022) and all procedures were carried out in accordance with the Declaration of Helsinki.

## Consent to participate

After detailed explanation of the study protocol, all subjects gave written informed consent. All subjects were insured and reimbursed for their participation.

## Data availability

Raw data will not be publicly available for data protection reasons. Processed data and custom code can be obtained from the corresponding author with a data sharing agreement, approved by the departments of legal affairs and data clearing of the Medical University of Vienna. The estimated relative cellular proportions and gene expression patterns are available from 10.5281/zenodo.23475383. Code related to the creation of maps and relative cellular proportions is available at https://github.com/NeuroimagingLabsMUV.

## Methods

### Marker gene definition

Cell-type marker genes were first identified based on the CellMarker ^47^ database, which is a curated resource of experimentally confirmed cell-type-specific markers. Starting from the original list of 1782 candidate markers, snRNA-Seq-measured gene expression (log2 of counts per million [CPM])^42^ was used to estimate the specificity of marker genes, quantified as the difference between expression in the target cell type and the highest expression among all other cell types. Genes with the specificity score > 2.0 were retained. To further refine the selection, we considered biological stability (abs(logFC) < 0.5) ^52^ and the correlation between snRNAseq expression ^42^ profiles and AHBA microarray probes to estimate voxel-wise transcriptomic maps (r > 0.4), prioritising markers with both robust biological expression and cross-modality consistency. Biological stability is indicated by the log2-fold-change (logFC) derived from the linear model that assesses the gene’s effect in relation to living or postmortem status ^52^.

### AHBA microarray treatment

Microarray expression data were obtained from the Allen Human Brain Atlas (AHBA). Parts of the data treatment were adapted from Gryglewski et al.^44^. Probes were first reannotated using the abagen toolbox^18, 34, 81^ and then filtered using a presence-above-background (PAC) criterion provided by the Allen Brain Institute, retaining only those that exceeded background signal in at least 1% of 3702 original samples, leaving us with 38595 probes covering 18076 gene expressions.

Sample coordinates (MNI space) underwent nonlinear spatial normalisation ^45^ in addition to the linear transformation provided by the AHBA. To mitigate hemispheric sampling imbalance, samples were mirrored across the sagittal plane. Samples were assigned to the cortex, subcortex, cerebellum, and brainstem based on the ontology annotation provided in AHBA. 20 white matter samples were excluded from analysis. Finally, 3493 mirrored samples in the cortex, 1770 in the subcortex, 1214 in the brainstem, and 731 in the cerebellum were retained.

To reduce inter-individual variability, gene expression values were mean-centred using region-wise averages computed from regions present in all donors. Regional mean expression values were computed using the Allen Brain Reference Atlas to support normalisation procedures. Regions with fewer than four samples per donor were excluded.

Probe selection was refined independently for each gene to maximise biological relevance and spatial robustness. First, probes with low inter-probe consistency (weighted pairwise Spearman’s rank correlation r<0.3) were excluded. The remaining probes were evaluated using a regional similarity measure (RSV), and probes whose removal increased RSV in at least 3 donors were discarded. In addition to the original probe selection performed by Gryglewski et al., probe expression profiles were correlated with corresponding snRNAseq-derived gene expression ^42^ to ensure cross-modality consistency. Only probes achieving at least 90% of the maximum correlation within a gene were retained.

### Voxel-wise mRNA expression estimation

Spatial gene expression maps were generated using variogram modelling and ordinary kriging. For each marker gene, empirical variograms were calculated from AHBA sample expression values to characterise spatial dependence, and parametric variogram models were fitted (Python v3.12.7, GSTools 1.7.0). Voxel-wise expression was then estimated using ordinary kriging based on spatially weighted neighbouring samples in MATLAB (R2018b, MathWorks, Natick, USA). A grey-matter mask used for spatial interpolation was derived from the probabilistic Harvard-Oxford atlas, as implemented in FSL ^46^, for the cerebral cortex, subcortex, and brainstem. Cerebellum was used as defined in the SUIT atlas ^82, 83^. The probabilistic maps of regions were thresholded at 10%. Overlapping voxels were assigned to the region of highest probability.

### Cell-type deconvolution using weighted ridge regression

Cell-type RNA contributions were estimated using weighted non-negative ridge regression modelling weighted voxel-wise marker gene expression (target matrix Y) as a linear combination of cell-type-specific transcriptional signatures derived from snRNAseq data (reference matrix B). The regression was solved by minimising the least-squares objective (Eq. 1) using projected gradient descent

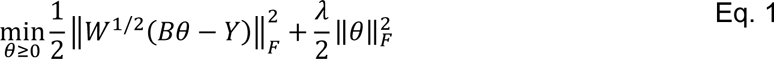

The step size of gradient descent was 1/L, where L is the Lipschitz constant defined by the largest eigenvalue of B^T^B+λ*I ^84^. The ridge parameter λ controls the strength of the penalty term on regression coefficients, controlling overfitting and stabilising the estimates in the presence of correlated predictors. The reference matrix B had 28 rows (one per marker gene) and three columns representing the cell types (neuronal, glial, vascular). Each element represented the mean CPM expression value (log2-transformed). To account for differences in total RNA abundance between cell types, the reference matrix was further adjusted using RNA-content weights derived from mean total UMI counts per nucleus ^42^. The reference matrix also included a dummy component column to account for unmodeled variation. The target matrix Y contained a voxel-wise representation of marker genes created in the previous step. These were further weighted by gene-specific weights (W) calculated as the product of a biological stability metric ^52^ and the correlation between snRNAseq expression profiles and AHBA microarray probes used to estimate voxel-wise transcriptomic maps, prioritising markers with both robust biological expression and cross-modality consistency. The resulting coefficients θ represent RNA-weighted cell-type contributions rather than direct cell counts. These coefficients are subsequently normalised to sum to 1 within each voxel, yielding relative contributions of neuronal (NEU), glial (GLI), and vascular (VAS) cell contribution. For a detailed explanation of individual steps, please refer to the supplementary material.

### Baseline glucose metabolism

To represent baseline glucose metabolism, we have used data from a previously published study^85^. Briefly, eighteen healthy volunteers (25.11±5.15 years; 12 females) were scanned with the radiotracer [^18^F]FDG on a large-axial-field-of-view PET/CT scanner (Biograph Vision Quadra, Siemens Healthineers, Erlangen, Germany). Each session comprised a 20-minute resting-state block with eyes open (EO) and 70 minutes with eyes closed (EC).

All participants underwent a standard medical examination done by a trained psychiatrist. At the initial screening visit, routine blood tests, electrocardiography, neurological tests, and the Structured Clinical Interview for DSM-IV were performed. All participants gave written informed consent, and all procedures were conducted in accordance with the Declaration of Helsinki. The study was approved by the ethics committee of the Medical University of Vienna (Ethics Number: 1642/2022)

The radioligand was administered as a bolus (816 ml/h for 1 minute) followed by a constant infusion (21 ml/h for 104 minutes). PET data were acquired in list mode and reconstructed post hoc into 1 s frames using Siemens’ e7 tools. An additional T1-weighted MPRAGE anatomical scan (echo-time/repetition-time=2.37/1800 ms, inversion-time=900 ms, acquisition-time=219 s, flip angle=8°, matrix size=288 × 288, 208 slices, voxel size=0.87 × 0.87 × 0.85 mm) was acquired on a Siemens Prisma 3T MR system (Siemens, Erlangen, Germany). This scan was used to normalise the PET images to the common stereotactic Montreal Neurological Institute space with SPM12 (Statistical Parametric Mapping, Release 12, Wellcome Centre for Human Neuroimaging, London, UK).

Quantification of CMRGlu was performed using the fPET toolbox ^86^. Venous blood samples were collected 4, 90 and 105 minutes after start of radiotracer injection. These samples, together with an image-derived input function from the descending aorta, were used as inputs to the Patlak plot to estimate CMRGlu. For the purposes of this work, we averaged across subjects to obtain mean CMRGlu maps for the EO and EC conditions.

### Mitochondrial metrics

Regional mitochondrial density maps were obtained from a publicly available dataset ^27^. The mitochondrial density was estimated using spatially resolved measurements of mitochondrial-related markers across the human brain and integrated into a regional map that reflects variation in mitochondrial abundance. The resulting mitochondrial density (MitoD) captures regional differences in mitochondrial content.

### Histological ground truth data

Representative distributions of neuronal and non-neuronal cells were obtained from published data ^48^. In particular, the proportions of neuronal and non-neuronal cells in the cortex, as well as the non-neuronal-to-neuronal-cell ratio, were estimated from histological data. Neurons and non-neuronal (NNE) cells were quantified using the isotropic fractionator on 101 coronal sections covering seven gross brain regions (prefrontal, parietal, temporal, dorsal, occipital, insula, V1) in three brains. Voxel-wise representations of mitochondrial density and CMRGlu were aligned with the dissections. Comparisons of the estimated relative cellular proportions or ratios (NEU_R_, GLI_R_ and NNE_R_/NEU_R_ ratio; NNE_R_=GLI_R_ + VAS_R_) were performed using Pearson correlation.

### Integration of cellular composition and mitochondrial density with glucose metabolism

For this analysis, we utilized a weighted linear regression model to elucidate the relationships among CMRGlu during eyes-closed and eyes-open condition, mitochondrial density, glia-to-neuron ratio (G2N). In the follow-up exploratory analysis, we replaced the G2N ratio with relative neuronal proportion or relative glial proportion. An interaction term between the cellular metric and mitochondrial density was included to evaluate both independent associations and potential modulation of cell-type effects by mitochondrial content. All variables were z-scored before analysis, and models were weighted by regional size to account for differences in anatomical sampling.

To further evaluate the involvement of cellular subtypes and mitochondria in the changes in glucose metabolism during eyes-open condition vs eyes-closed, we assessed the relation of G2N and MitoD to the percent signal change in CMRGlu (PSC_CMRGlu_ = [EO/EC - 1] × 100). Here we also introduce a quadratic term to capture the non-linear change in cellular involvement. In the subsequent exploratory analysis, we have replaced G2N by NEU_R_ and GLI_R_. Likewise, G2N was replaced by the CUX1, CUX2, FEZF2, and BCL11B expression patterns to investigate the relationship of axonal calibre to PSC_CMRGlu_.

Post hoc analyses were performed using Pearson correlations between the covariates and CMRGlu, corrected for the respective variable, as well as for mitochondrial density and cell-type proportions. Each variable was represented by 105 cortical regions defined in the Brainetome atlas ^87^. The hierarchical structure of the Brainetome atlas allowed the subsequent separation of the analysis into six major lobes: frontal, temporal, parietal, insular, limbic, and occipital. To account for spatial autocorrelation arising from the spatial dependence between regions ^88^, we performed a spin test with Vásquez-Rodríguez’s ^89^ spatial permutation model to obtain spatially corrected p-values (p_spin_).

## References

1. Dienel, G.A. Brain Glucose Metabolism: Integration of Energetics with Function. Physiological Reviews 99, 949–1045 (2019).

2. Herculano-Houzel, S. Scaling of Brain Metabolism with a Fixed Energy Budget per Neuron: Implications for Neuronal Activity, Plasticity and Evolution. PLoS ONE 6, e17514 (2011).

3. McKenna, M.C., Dienel, G.A., Sonnewald, U., Waagepetersen, H.S. & Schousboe, A. Chapter 11 - Energy Metabolism of the Brain. in Basic Neurochemistry (Eighth Edition) (ed. S.T. Brady, G.J. Siegel, R.W. Albers & D.L. Price) 200–231 (Academic Press, New York, 2012).

4. Raichle, M.E. & Gusnard, D.A. Appraising the brain’s energy budget. Proc. Natl. Acad. Sci. U.S.A. 99, 10237–10239 (2002).

5. Attwell, D. & Laughlin, S.B. An Energy Budget for Signaling in the Grey Matter of the Brain. J Cereb Blood Flow Metab 21, 1133–1145 (2001).

6. Harris, Julia J., Jolivet, R. & Attwell, D. Synaptic Energy Use and Supply. Neuron 75, 762–777 (2012).

7. Pourmajidian, M., Hansen, J.Y., Shafiei, G., Misic, B. & Dagher, A. Five energy metabolism pathways show distinct regional distributions and lifespan trajectories in the human brain. PLoS Biol 24, e3003619 (2026).

8. Mergenthaler, P., Lindauer, U., Dienel, G.A. & Meisel, A. Sugar for the brain: the role of glucose in physiological and pathological brain function. Trends in Neurosciences 36, 587–597 (2013).

9. Huang, S.C., et al. Noninvasive determination of local cerebral metabolic rate of glucose in man. American Journal of Physiology-Endocrinology and Metabolism 238, E69–E82 (1980).

10. Wadsak, W. & Mitterhauser, M. Basics and principles of radiopharmaceuticals for PET/CT. European Journal of Radiology 73, 461–469 (2010).

11. Sokoloff, L. Localization of Functional Activity in the Central Nervous System by Measurement of Glucose Utilization with Radioactive Deoxyglucose. J Cereb Blood Flow Metab 1, 7–36 (1981).

12. Hahn, A., et al. High-temporal resolution functional PET/MRI reveals coupling between human metabolic and hemodynamic brain response. Eur J Nucl Med Mol Imaging 51, 1310–1322 (2024).

13. Jamadar, S.D., et al. Simultaneous task-based BOLD-fMRI and [18-F] FDG functional PET for measurement of neuronal metabolism in the human visual cortex. NeuroImage 189, 258–266 (2019).

14. Villien, M., et al. Dynamic functional imaging of brain glucose utilization using fPET-FDG. NeuroImage 100, 192–199 (2014).

15. Bélanger, M., Allaman, I. & Magistretti, Pierre J. Brain Energy Metabolism: Focus on Astrocyte-Neuron Metabolic Cooperation. Cell Metabolism 14, 724–738 (2011).

16. Xiang, X., et al. Microglial activation states drive glucose uptake and FDG-PET alterations in neurodegenerative diseases. Sci. Transl. Med. 13, eabe5640 (2021).

17. Zimmer, E.R., et al. [18F]FDG PET signal is driven by astroglial glutamate transport. Nature Neuroscience 20, 393–395 (2017).

18. Arnatkevičiūtė, A., Fulcher, B.D. & Fornito, A. A practical guide to linking brain-wide gene expression and neuroimaging data. NeuroImage 189, 353–367 (2019).

19. Hansen, J.Y. & Misic, B. Integrating and interpreting brain maps. Trends in Neurosciences 48, 594–607 (2025).

20. Monzel, A.S., Enríquez, J.A. & Picard, M. Multifaceted mitochondria: moving mitochondrial science beyond function and dysfunction. Nat Metab 5, 546–562 (2023).

21. Hara, Y., et al. Presynaptic mitochondrial morphology in monkey prefrontal cortex correlates with working memory and is improved with estrogen treatment. Proc. Natl. Acad. Sci. U.S.A. 111, 486-491 (2014).

22. Sharpley, Mark S., et al. Heteroplasmy of Mouse mtDNA Is Genetically Unstable and Results in Altered Behavior and Cognition. Cell 151, 333–343 (2012).

23. Picard, M. & Shirihai, O.S. Mitochondrial signal transduction. Cell Metabolism 34, 1620–1653 (2022).

24. Styr, B., et al. Mitochondrial Regulation of the Hippocampal Firing Rate Set Point and Seizure Susceptibility. Neuron 102, 1009–1024.e1008 (2019).

25. Kwon, S.-K., et al. LKB1 Regulates Mitochondria-Dependent Presynaptic Calcium Clearance and Neurotransmitter Release Properties at Excitatory Synapses along Cortical Axons. PLoS Biol 14, e1002516 (2016).

26. Erecińska, M. & Silver, I.A. ATP and Brain Function. J Cereb Blood Flow Metab 9, 2–19 (1989).

27. Mosharov, E.V., et al. A human brain map of mitochondrial respiratory capacity and diversity. Nature 641, 749–758 (2025).

28. Sager, G., et al. Spatial and morphological organization of mitochondria in neurons across a connectome. Science 391, eads6674 (2026).

29. Gebara, E., et al. Mitofusin-2 in the Nucleus Accumbens Regulates Anxiety and Depression-like Behaviors Through Mitochondrial and Neuronal Actions. Biological Psychiatry 89, 1033–1044 (2021).

30. Hollis, F., et al. Mitochondrial function in the brain links anxiety with social subordination. Proc. Natl. Acad. Sci. U.S.A. 112, 15486-15491 (2015).

31. Joshi, A.U., et al. Fragmented mitochondria released from microglia trigger A1 astrocytic response and propagate inflammatory neurodegeneration. Nature Neuroscience 22, 1635–1648 (2019).

32. Lin, M.-m., Liu, N., Qin, Z.-h. & Wang, Y. Mitochondrial-derived damage-associated molecular patterns amplify neuroinflammation in neurodegenerative diseases. Acta Pharmacol Sin 43, 2439–2447 (2022).

33. Arnatkeviciute, A., Markello, R.D., Fulcher, B.D., Misic, B. & Fornito, A. Toward Best Practices for Imaging Transcriptomics of the Human Brain. Biological Psychiatry 93, 391–404 (2023).

34. Hawrylycz, M.J., et al. An anatomically comprehensive atlas of the adult human brain transcriptome. Nature 489, 391–399 (2012).

35. Chen, X., et al. A brain cell atlas integrating single-cell transcriptomes across human brain regions. Nat Med 30, 2679–2691 (2024).

36. Shin, J., et al. Cell-Specific Gene-Expression Profiles and Cortical Thickness in the Human Brain. Cerebral Cortex 28, 3267–3277 (2018).

37. Van Aalst, J., et al. In vivo synaptic density relates to glucose metabolism at rest in healthy subjects, but is strongly modulated by regional differences. J Cereb Blood Flow Metab 41, 1978–1987 (2021).

38. Markello, R.D., et al. neuromaps: structural and functional interpretation of brain maps. Nat Methods 19, 1472–1479 (2022).

39. Ng, B., et al. Integration across biophysical scales identifies molecular and cellular correlates of person-to-person variability in human brain connectivity. Nature Neuroscience 27, 2240–2252 (2024).

40. Hahn, A., et al. [18F]FDG Revisited: A New Perspective on the Temporal Dynamics of Brain Glucose Metabolism. (Medicine and Pharmacology, 2025).

41. Seidlitz, J., et al. Transcriptomic and cellular decoding of regional brain vulnerability to neurogenetic disorders. Nat Commun 11, 3358 (2020).

42. Siletti, K., et al. Transcriptomic diversity of cell types across the adult human brain. Science 382, eadd7046 (2023).

43. Zhang, X.-H., et al. The cell-type underpinnings of the human functional cortical connectome. Nature Neuroscience 28, 150–160 (2025).

44. Gryglewski, G., et al. Spatial analysis and high resolution mapping of the human whole-brain transcriptome for integrative analysis in neuroimaging. NeuroImage 176, 259–267 (2018).

45. Murgaš, M., Michethaler, P., Reed, M.B., Gryglewski, G. & Lanzenberger, R. Correlation of receptor density and mRNA expression patterns in the human cerebral cortex. NeuroImage 256, 119214 (2022).

46. Jenkinson, M., Beckmann, C.F., Behrens, T.E., Woolrich, M.W. & Smith, S.M. FSL. NeuroImage 62, 782–790 (2012).

47. Zhang, X., et al. CellMarker: a manually curated resource of cell markers in human and mouse. Nucleic Acids Research 47, D721–D728 (2019).

48. Ribeiro, P.F.M., et al. The human cerebral cortex is neither one nor many: neuronal distribution reveals two quantitatively different zones in the gray matter, three in the white matter, and explains local variations in cortical folding. Front. Neuroanat. 7, 28 (2013).

49. Sherwood, C.C., et al. Evolution of increased glia-neuron ratios in the human frontal cortex. Proc Natl Acad Sci U S A 103, 13606–13611 (2006).

50. Von Bartheld, C.S., Bahney, J. & Herculano-Houzel, S. The search for true numbers of neurons and glial cells in the human brain: A review of 150 years of cell counting. J of Comparative Neurology 524, 3865–3895 (2016).

51. Hoerl, A.E. & Kennard, R.W. Ridge Regression: Biased Estimation for Nonorthogonal Problems. Technometrics 12, 55–67 (1970).

52. Liharska, L.E., et al. A study of gene expression in the living human brain. (Genetic and Genomic Medicine, 2023).

53. Langseth, C.M., et al. Comprehensive in situ mapping of human cortical transcriptomic cell types. Communications Biology 4, 998 (2021).

54. Azevedo, F.A.C., et al. Equal numbers of neuronal and nonneuronal cells make the human brain an isometrically scaled-up primate brain. J of Comparative Neurology 513, 532–541 (2009).

55. Magistretti, Pierre J. & Allaman, I. A Cellular Perspective on Brain Energy Metabolism and Functional Imaging. Neuron 86, 883–901 (2015).

56. Theriault, J.E., et al. A functional account of stimulation-based aerobic glycolysis and its role in interpreting BOLD signal intensity increases in neuroimaging experiments. Neuroscience & Biobehavioral Reviews 153, 105373 (2023).

57. Corbetta, M. & Shulman, G.L. Control of goal-directed and stimulus-driven attention in the brain. Nat Rev Neurosci 3, 201–215 (2002).

58. Sengupta, B., Faisal, A.A., Laughlin, S.B. & Niven, J.E. The Effect of Cell Size and Channel Density on Neuronal Information Encoding and Energy Efficiency. J Cereb Blood Flow Metab 33, 1465–1473 (2013).

59. Pak, V., et al. Distinctive whole-brain cell types predict tissue damage patterns in thirteen neurodegenerative conditions. Elife 12, RP89368 (2024).

60. Sutton, G.J., et al. Comprehensive evaluation of deconvolution methods for human brain gene expression. Nat Commun 13, 1358 (2022).

61. Garcia-Marin, V., Kelly, J.G. & Hawken, M.J. Neuronal composition of processing modules in human V1: laminar density for neuronal and non-neuronal populations and a comparison with macaque. Cerebral Cortex 34, bhad512 (2024).

62. Herculano-Houzel, S. The glia/neuron ratio: How it varies uniformly across brain structures and species and what that means for brain physiology and evolution. Glia 62, 1377–1391 (2014).

63. Howarth, C., Gleeson, P. & Attwell, D. Updated energy budgets for neural computation in the neocortex and cerebellum. J Cereb Blood Flow Metab 32, 1222–1232 (2012).

64. Engl, E. & Attwell, D. Non-signalling energy use in the brain. The Journal of Physiology 593, 3417–3429 (2015).

65. Amunts, K. & Zilles, K. Architectonic Mapping of the Human Brain beyond Brodmann. Neuron 88, 1086–1107 (2015).

66. Uludağ, K., et al. Coupling of cerebral blood flow and oxygen consumption during physiological activation and deactivation measured with fMRI. NeuroImage 23, 148–155 (2004).

67. Brewer, G.J., et al. Neuron network activity scales exponentially with synapse density. J Neural Eng 6, 014001 (2009).

68. Raichle, M.E. & Mintun, M.A. Brain work and brain imaging. Annu Rev Neurosci 29, 449–476 (2006).

69. Bednařík, P., et al. Neurochemical and BOLD Responses during Neuronal Activation Measured in the Human Visual Cortex at 7 Tesla. J Cereb Blood Flow Metab 35, 601–610 (2015).

70. Díaz-García, C.M., et al. Neuronal Stimulation Triggers Neuronal Glycolysis and Not Lactate Uptake. Cell Metabolism 26, 361–374.e364 (2017).

71. Fox, P.T. & Raichle, M.E. Focal physiological uncoupling of cerebral blood flow and oxidative metabolism during somatosensory stimulation in human subjects. Proc. Natl. Acad. Sci. U.S.A. 83, 1140-1144 (1986).

72. Fox, P.T., Raichle, M.E., Mintun, M.A. & Dence, C. Nonoxidative glucose consumption during focal physiologic neural activity. Science 241, 462–464 (1988).

73. Cubelos, B., et al. Cux1 and Cux2 Regulate Dendritic Branching, Spine Morphology, and Synapses of the Upper Layer Neurons of the Cortex. Neuron 66, 523–535 (2010).

74. Nieto, M., et al. Expression of Cux-1 and Cux-2 in the subventricular zone and upper layers II–IV of the cerebral cortex. J of Comparative Neurology 479, 168–180 (2004).

75. Arlotta, P., et al. Neuronal Subtype-Specific Genes that Control Corticospinal Motor Neuron Development In Vivo. Neuron 45, 207–221 (2005).

76. Molyneaux, B.J., Arlotta, P., Hirata, T., Hibi, M. & Macklis, J.D. Fezl Is Required for the Birth and Specification of Corticospinal Motor Neurons. Neuron 47, 817–831 (2005).

77. Firmin, L., et al. Axon diameters and conduction velocities in the macaque pyramidal tract. Journal of Neurophysiology 112, 1229–1240 (2014).

78. Innocenti, G.M., et al. Diversity of Cortico-descending Projections: Histological and Diffusion MRI Characterization in the Monkey. Cerebral Cortex 29, 788–801 (2019).

79. Pascual-García, M., et al. Morphological correlates of pyramidal cell axonal myelination in mouse and human neocortex. Cerebral Cortex 34, bhae147 (2024).

80. Selvaggi, P., Rizzo, G., Mehta, M.A., Turkheimer, F.E. & Veronese, M. Integration of human whole-brain transcriptome and neuroimaging data: Practical considerations of current available methods. Journal of Neuroscience Methods 355, 109128 (2021).

81. Markello, R.D., et al. Standardizing workflows in imaging transcriptomics with the abagen toolbox. Elife 10, e72129 (2021).

82. Diedrichsen, J., Balsters, J.H., Flavell, J., Cussans, E. & Ramnani, N. A probabilistic MR atlas of the human cerebellum. NeuroImage 46, 39–46 (2009).

83. Diedrichsen, J. A spatially unbiased atlas template of the human cerebellum. NeuroImage 33, 127–138 (2006).

84. Beck, A. & Teboulle, M. A Fast Iterative Shrinkage-Thresholding Algorithm for Linear Inverse Problems. SIAM Journal on Imaging Sciences 2, 183–202 (2009).

85. Stöhrmann, P., et al. Small but systematic bias introduced by EEG electrodes in PET imaging. medRxiv, 2026.2008.2012.26360268 (2026).

86. Hahn, A., et al. A unified approach for identifying PET-based neuronal activation and molecular connectivity with the functional PET toolbox. J Cereb Blood Flow Metab 46, 558–568 (2026).

87. Fan, L., et al. The Human Brainnetome Atlas: A New Brain Atlas Based on Connectional Architecture. Cereb Cortex 26, 3508–3526 (2016).

88. Markello, R.D. & Misic, B. Comparing spatial null models for brain maps. NeuroImage 236, 118052 (2021).

89. Vázquez-Rodríguez, B., et al. Gradients of structure–function tethering across neocortex. Proc. Natl. Acad. Sci. U.S.A. 116, 21219-21227 (2019).

