## Supplementary material for "Cellular and mitochondrial correlates of regional glucose metabolism in the human cortex"

### Supplementary figures

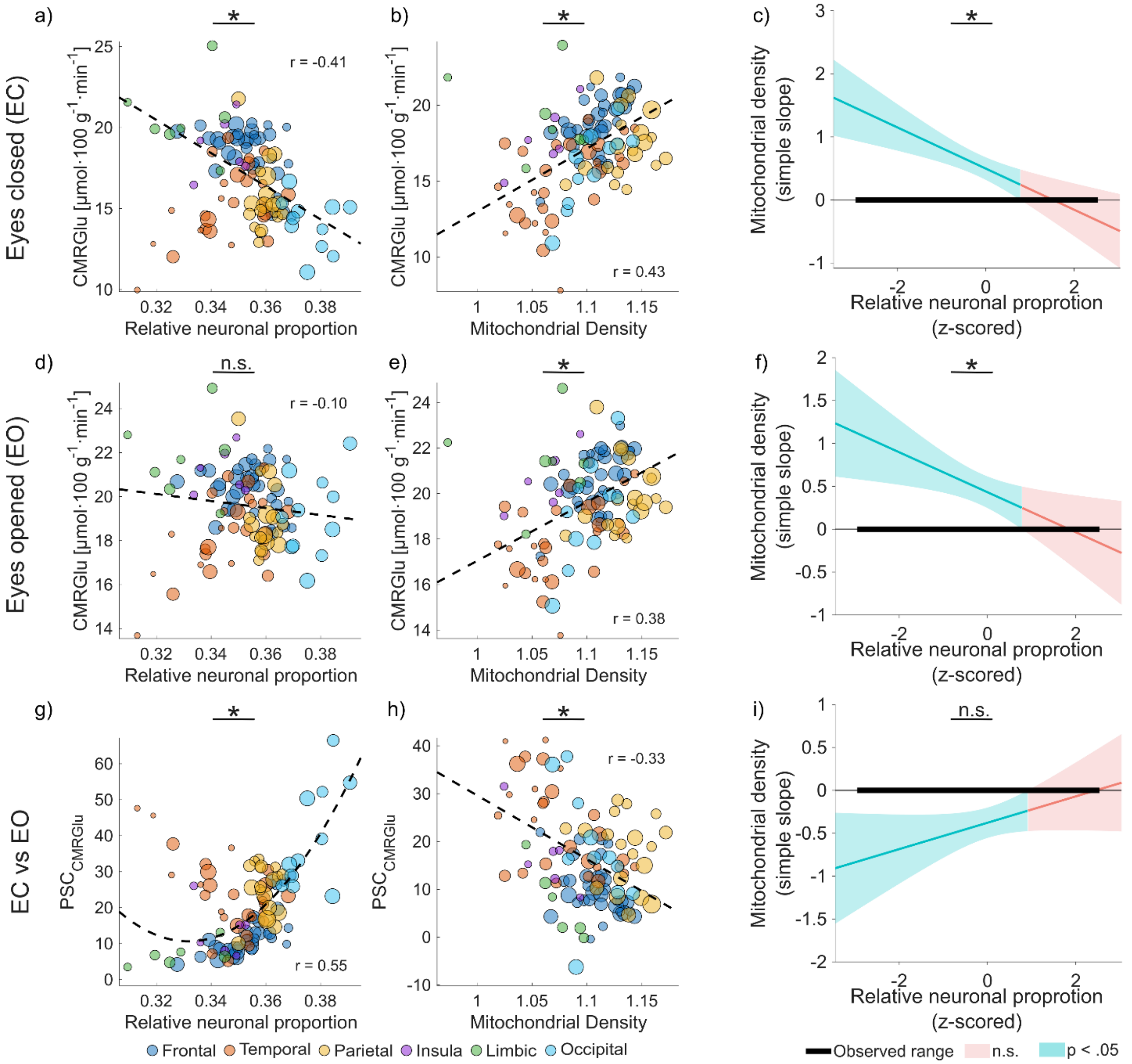


Suppl Figure 1: M**odel integrating baseline glucose metabolism, mitochondrial density and relative neuronal proportion.** Model investigating the impact of relative neuronal proportion ($NEU_R$) and mitochondrial density (MitoD), on glucose metabolism (CMRGlu). Resulting relationships are shown in the form of a colour-labelled scatter plot. The size of the marker represents the size of the regions used in weighted linear regression as a scaling factor, and the colour represents marker association with gross cortical regions - frontal (dark blue), temporal (red), parietal (yellow), insular (violet), limbic (green), and occipital (cyan) lobes. Observed associations between resting-state CMRGlu during eyes-closed conditions ($CMRGlu_{EC}$) and (a) the $NEU_R$ and (b) mitochondrial density (MitoD) are in the first row. Association of CMRGlu during the eyes-open ($CMRGlu_{EO}$) and d) $NEU_R$ and e) MitoD is in the middle row of the figure. Association of the MitoD and $NEU_R$ ratio with the per cent signal change ($PSC_{CMRGlu} = CMRGlu_{EO}/CMRGlu_{EC} - 1$) is shown in panels g) and h), respectively. This model includes a quadratic term to compensate for nonlinear change in the CMRGlu. The correlation coefficient represents the relationship between the $NEU_R$ and $PSC_{CMRGlu}$ (excluding the nonlinear term). The interaction effects between the $NEU_R$ and MitoD during c) eyes closed, f) eyes opened and i) $PSC_{CMRGlu}$ are represented in the Johnson-Neyman plot. Johnson-Neyman plot shows the conditional slopes of MitoD as a function of $NEU_R$. The model including a quadratic term shows the total slope of the $NEU_R$ as a function of MitoD, accounting for a non-linear term. Solid lines represent the model-estimated simple slope at each value of the moderator; shaded bands represent the corresponding 95% confidence interval. Teal shading indicates values of the moderator at which the simple slope is statistically significant ($p < 0.05$); pink shading indicates the slope is not significantly different from zero. The thick black horizontal bar denotes the observed range of the moderator variable in the sample. Statistically significant post-hoc test are marked with * and non-significant as n.s..

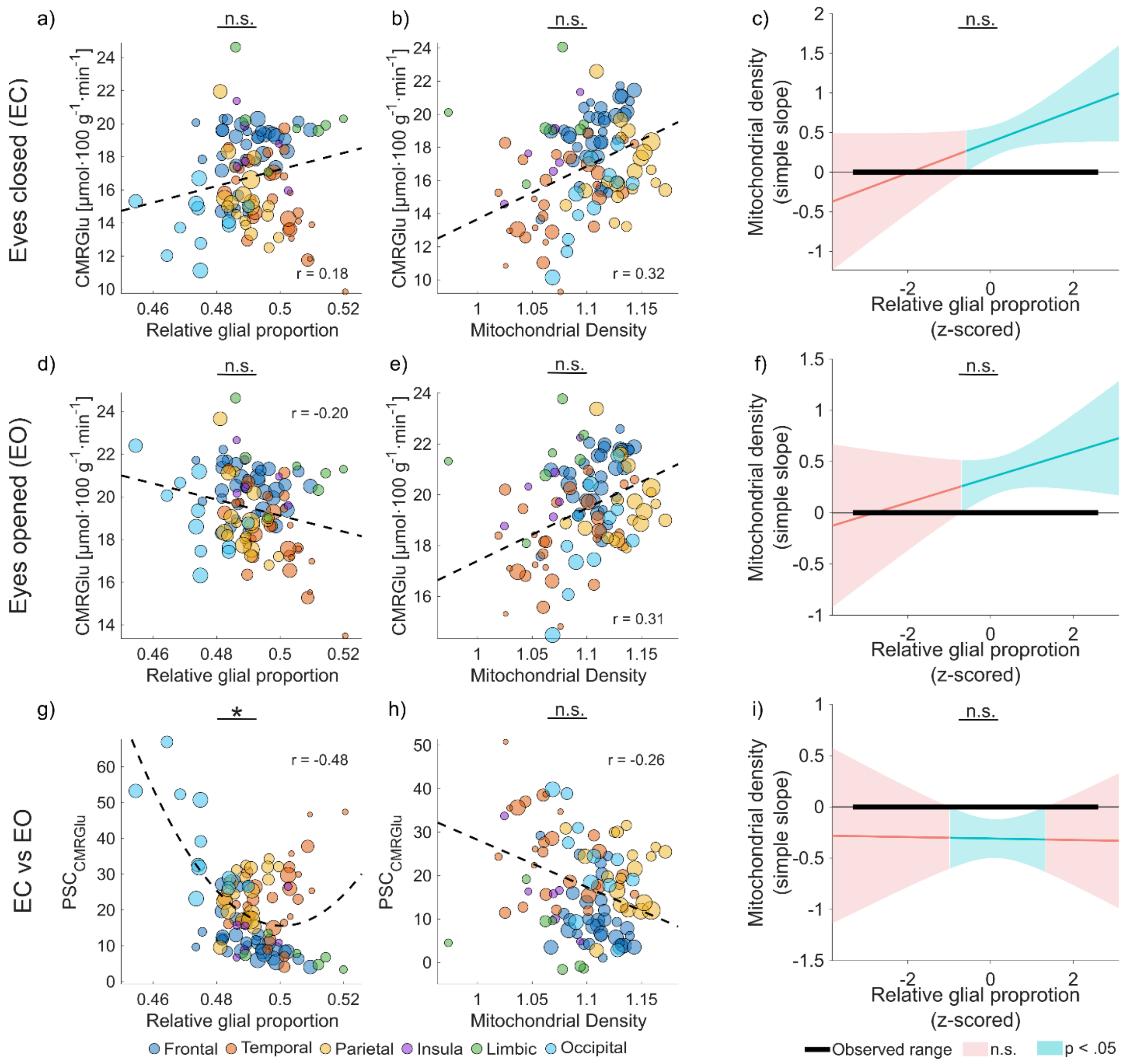


Suppl Figure 2: M**odel integrating baseline glucose metabolism, mitochondrial density and relative glial proportion.** Model investigating the impact of relative glial proportion ($GLI_R$) and mitochondrial density (MitoD), on glucose metabolism (CMRGlu). Resulting relationships are shown in the form of a colour-labelled scatter plot. The size of the marker represents the size of the regions used in weighted linear regression as a scaling factor, and the colour represents marker association with gross cortical regions - frontal (dark blue), temporal (red), parietal (yellow), insular (violet), limbic (green), and occipital (cyan) lobes. Observed associations between resting-state CMRGlu during eyes-closed conditions ($CMRGlu_{EC}$) and (a) the $GLI_R$ and (b) mitochondrial density (MitoD) are in the first row. Association of CMRGlu during the eyes-open ($CMRGlu_{EO}$) and d) $GLI_R$ and e) MitoD is in the middle row of the figure. Association of the MitoD and $GLI_R$ ratio with the per cent signal change ($PSC_{CMRGlu} = CMRGlu_{EO}/CMRGlu_{EC} - 1$) is shown in panels g) and h), respectively. This model includes a quadratic term to compensate for nonlinear change in the CMRGlu. The correlation coefficient represents the relationship between the $GLI_R$ and $PSC_{CMRGlu}$ (excluding the nonlinear term). The interaction effects between the $GLI_R$ and MitoD during c) eyes closed, f) eyes opened and i) $PSC_{CMRGlu}$ are represented in the Johnson-Neyman plot. Johnson-Neyman plot shows the conditional slopes of MitoD as a function of $GLI_R$. The model including a quadratic term shows the total slope of the $GLI_R$ as a function of MitoD, accounting for a non-linear term. Solid lines represent the model-estimated simple slope at each value of the moderator; shaded bands represent the corresponding 95% confidence interval. Teal shading indicates values of the moderator at which the simple slope is statistically significant ($p < 0.05$); pink shading indicates the slope is not significantly different from zero. The thick black horizontal bar denotes the observed range of the moderator variable in the sample. Statistically significant post-hoc test are marked with * and non-significant as n.s..

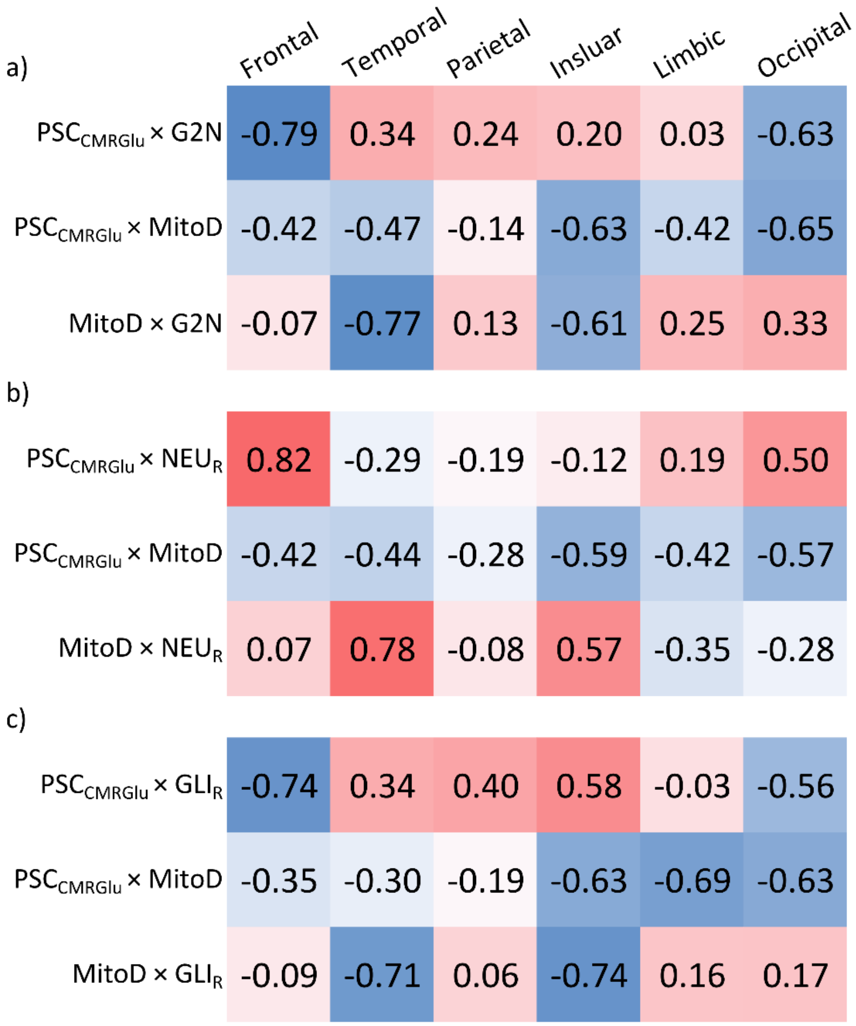


Suppl Figure 3: Heatmaps showing the association (weighted Spearman's ranked correlation coefficient) between $PSC_{CMRGlu}$, mitochondrial density (MitoD) and a) glia-to-neuron ratio, b) relative neuronal proportion ($NEU_R$) and c) relative glial proportion ($GLI_R$) in the frontal, temporal, parietal, insular, limbic and occipital lobe. Prior to correlation analysis, each $PSC_{CMRGlu}$ is corrected for the respective variable.

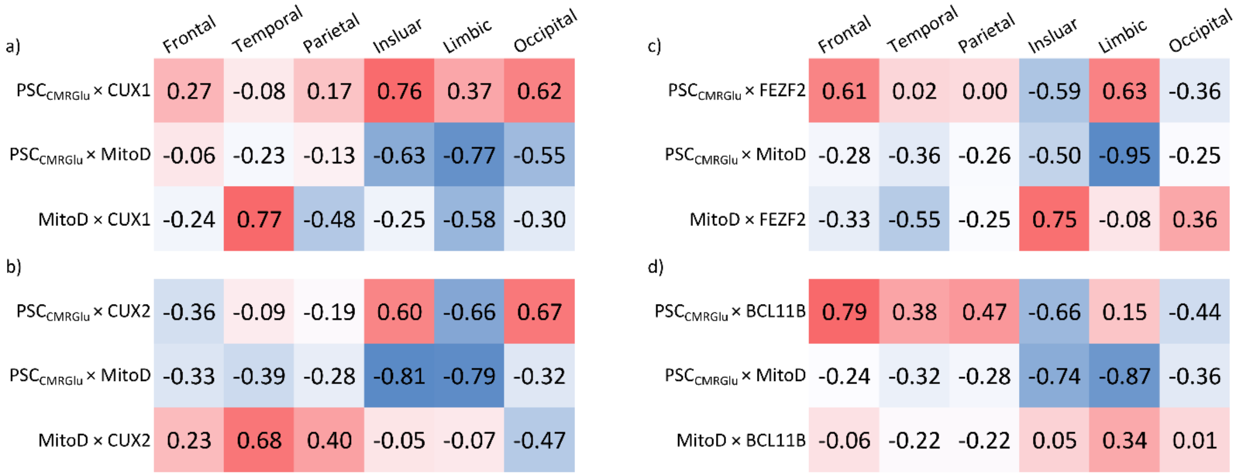


Suppl Figure 4: Heatmaps showing the association (weighted Spearman's ranked correlation coefficient) between $PSC_{CMRGlu}$, mitochondrial density (MitoD) and thin-axonal caliber marker genes a) Cut-like Homeobox 1 (CUX1) and b) Cut-like Homeobox 2 (CUX2) and thick-axonal calibre marker genes c) Fez family zinc finger 2 (FEZF2) and d) BCL11 transcription factor B (BCL11B), in the frontal, temporal, parietal, insular, limbic and occipital lobe. Prior to correlation analysis, each $PSC_{CMRGlu}$ is corrected for the respective variable.

Supplementary tables

| Neuronal cells | | Glial cells | | Vascular cells | |
|---|---|---|---|---|---|
| Gene Symbol | Gene Name | Gene Symbol | Gene Name | Gene Symbol | Gene Name |
| NRG1 | neuregulin 1 | GFAP | glial fibrillary acidic protein | EBF1 | early B cell factor 1 |
| RBFOX3 | RNA binding fox-1 homolog 3 | ITGAM | integrin subunit alpha M | COL1A1 | collagen type I alpha 1 chain |
| GAD2 | glutamate decarboxylase 2 | MOG | myelin oligodendrocyte glycoprotein | FOXF2 | forkhead box F2 |
| GABRA1 | gamma-aminobutyric acid type A receptor subunit alpha1 | S100A1 | S100 calcium binding protein A1 | OCLN | occludin |
| BCL11B | BCL11 transcription factor B | SLC1A3 | solute carrier family 1 member 3 | NOTCH3 | notch receptor 3 |
| EPHA7 | EPH receptor A7 | AQP4 | aquaporin 4 | COL1A2 | collagen type I alpha 2 chain |
| CNTN4 | contactin 4 | C1QB | complement C1q B chain | MYL9 | myosin light chain 9 |
| SLC17A6 | solute carrier family 17 member 6 | MAG | myelin associated glycoprotein | DCN | decorin |
| DLX1 | distal-less homeobox 1 | PDGFRA | platelet derived growth factor receptor alpha | VCAM1 | vascular cell adhesion molecule 1 |
| NPY | neuropeptide Y | | | | |

Suppl Table 1: List of selected marker genes for the neuronal (left), glial (middle) and vascular (right) cells.

| | **Estimate** | **SE** | **tStat** | **$p_{spin}$** |
|---|---|---|---|---|
| **$NEU_R$** | 0.629 | 0.091 | 6.918 | <0.05 |
| **$NEU_R^2$** | 0.217 | 0.059 | 3.699 | <0.05 |
| **MitoD** | -0.378 | 0.089 | -4.236 | <0.05 |
| **CT:M** | 0.154 | 0.090 | 1.704 | 0.27 |

Suppl Table 2: Results of the weighted linear regression integrating relative neuronal proportion ($NEU_R$), mitochondrial density (MitoD) and change in the glucose metabolism elicited by the change in functional state ($PSC_{CMRGlu}$)

| | **Estimate** | **SE** | **tStat** | **$p_{spin}$** |
|---|---|---|---|---|
| **$GLI_R$** | -0.364 | 0.100 | -3.647 | <0.05 |
| **$GLI_R^2$** | 0.221 | 0.060 | 3.688 | 0.09 |
| **MitoD** | -0.308 | 0.096 | -3.214 | 0.11 |
| **CT:M** | -0.007 | 0.107 | -0.067 | 0.96 |

Suppl Table 3: Results of the weighted linear regression integrating relative glial proportion ($GLI_R$), mitochondrial density (MitoD) and change in the glucose metabolism elicited by the change in functional state ($PSC_{CMRGlu}$)
